# Translation elongation as a rate limiting step of protein production

**DOI:** 10.1101/2023.11.27.568910

**Authors:** Elijah F Lyons, Lou C Devanneaux, Ryan Y Muller, Anna V Freitas, Zuriah A Meacham, Maria V McSharry, Van N Trinh, Anna J Rogers, Nicholas T Ingolia, Liana F Lareau

## Abstract

The impact of synonymous codon choice on protein output has important implications for understanding endogenous gene expression and design of synthetic mRNAs. Synonymous codons are decoded at different speeds, but simple models predict that this should not drive protein output. Instead, translation initiation should be the rate limiting step for production of protein per mRNA, with little impact of codon choice. Previously, we used a neural network model to design a series of synonymous fluorescent reporters and showed that their protein output in yeast spanned a seven-fold range corresponding to their predicted translation elongation speed. Here, we show that this effect is not due primarily to the established impact of slow elongation on mRNA stability, but rather, that slow elongation further decreases the number of proteins made per mRNA. We combine simulations and careful experiments on fluorescent reporters to show that translation is limited on non-optimally encoded transcripts. Using a genome-wide CRISPRi screen, we find that impairing translation initiation attenuates the impact of slow elongation, showing a dynamic balance between rate limiting steps of protein production. Our results show that codon choice can directly limit protein production across the full range of endogenous variability in codon usage.

## Introduction

The genetic code is elegant yet redundant, with 20 amino acids specified by 61 codons. The synonymous codon options are not exactly equivalent. Genomes have evolved to prefer some codons over others, and the differences between synonymous codons have real impact. As a ribosome moves along an mRNA, it spends more time translating some codons than others, because some tRNAs are less available or interact less favorably with the translation machinery. High resolution methods have allowed a close look at translation, revealing surprising consequences of codon choice for mRNA stability, protein output, and proper co-translational protein folding. Decoding the impact of synonymous codon usage is key to a more complete understanding of the output of the genome as well as better models for the design of synthetic mRNAs such as those used in vaccines.

It was not initially clear that codon choice or the speed of translation elongation would generally affect protein output. In a simple model where ribosomes initiate infrequently and elongation proceeds at a rate that is fast relative to initiation, the total number of proteins made per mRNA per time would not depend on the speed of elongation (Plotkin and Kudla 2011; Shah et al. 2013; Erdmann-Pham et al. 2020). Each ribosome that initiates would successfully reach the end of the reading frame to make one protein and the total protein production per mRNA per time would depend only on translation initiation. Translation initiation was recently measured to take 30 seconds in an *in vitro* yeast system (Wang et al. 2022) while translation elongation proceeds at 6 amino acids per second in mammalian cells (Ingolia et al. 2011), suggesting that elongation is rarely able to limit output (Plotkin and Kudla 2011; Shah et al. 2013; Riba et al. 2019; Erdmann-Pham et al. 2020).

However, groundbreaking results have established solidly that codon choice does affect protein output. Most notably, slow translation can destabilize mRNAs and thereby reduce total protein production, a process termed codon optimality mediated decay (Presnyak et al. 2015; Bazzini et al. 2016; Narula et al. 2019; Q. Wu et al. 2019; Bae and Coller 2022). Slow decoding causes a ribosome to spend more time without the proper tRNA engaged, and ribosomes in this conformation are more likely to recruit mRNA decay machinery, linking translation elongation speed to mRNA stability and indirectly limiting protein output by reducing mRNA abundance (Buschauer et al. 2020). In this way, translation elongation becomes limiting for protein synthesis without violating the traffic flow assumptions described above.

Variability in translation elongation speed can be captured empirically with ribosome profiling, a high throughput method that captures the positions of individual ribosomes and shows their distribution along mRNAs (Ingolia et al. 2009). The experiment is more likely to sample ribosomes at positions where they spend more time, i.e., positions that are decoded slowly. The count of ribosome footprints at a position thus reflects the dwell time of ribosomes decoding that position, when considered relative to the average count of footprints on that gene to control for differences in transcription and overall translation (Lareau et al. 2014). Thus, computational analysis of ribosome profiling data can reveal the progression of ribosomes along transcripts with remarkable resolution.

In our previous work, we made use of ribosome profiling data and machine learning to establish the connection between codon identity, ribosome speed, and protein output (Tunney et al. 2018). We used a neural network to learn the relationship between coding sequence and ribosome dwell time for the yeast *S. cerevisiae*, predicting the empirical ribosome footprint counts at each position based on the mRNA sequence in and near the A site of the ribosome. The predicted footprint counts can be interpreted as dwell times and summed to predict the total time to decode a sequence. We went on to ask if this total elongation time was also a good predictor of protein output, testing not only the aptness of our model but also the underlying question of how elongation speed affected protein output. We used our model to design six synonymous encodings of a fluorescent protein, citrine, with predicted decoding times spanning the range of endogenous yeast genes. When we expressed these reporters in yeast, we found a striking linear correspondence between the predicted decoding time and the protein output for each variant. Importantly, the translation efficiency, or proteins produced per mRNA in a given amount of time, also tracked the predicted decoding time. The difference in output from our reporters was only partially due to mRNA decay effects; unexpectedly, translation elongation speed also seemed to limit the rate of protein output more directly.

By a simple model of ribosome flow along transcripts, our synonymous constructs should each produce roughly the same protein per mRNA per time; additional processes must contribute to create a difference in translation efficiency. This observation motivated the current study to identify the scenarios under which slow codons limit protein output. Fundamentally, to produce less protein per mRNA, slow elongation must either cause fewer ribosomes to initiate, cause fewer ribosomes to terminate successfully, or produce less-stable proteins, and these outcomes may depend on the specific parameters of a transcript’s translation.

Translation elongation could conceivably be rate limiting through three potential paths. If elongation is very slow relative to initiation, ribosome traffic jams could back up to the start codon and occlude it, preventing initiation until the jam is cleared (Plotkin and Kudla 2011; Shah et al. 2013; Chu et al. 2014; Tuller et al. 2010; Riba et al. 2019; Verma et al. 2019; Erdmann-Pham et al. 2020). This would not require any active mechanism beyond the physical process of ribosomes moving slowly along an mRNA. Active mechanisms could also be implicated, perhaps by preventing ribosomes from reaching the end of the coding region. Acute ribosome stalls and collisions in both yeast and mammalian cells cause decreased protein output via processes like no-go decay or non-stop decay that couple abortive termination and mRNA degradation, followed by nascent chain degradation through ribosome-associated quality control (Shoemaker and Green 2012; Brandman and Hegde 2016; Meydan and Guydosh 2021; Brandman et al. 2012; Veltri et al. 2022). Similar mechanisms could decrease the protein output from slowly decoded yet non-defective transcripts. Finally, some feedback mechanism could sense slow elongation and limit initiation. A pathway in which collided ribosomes inhibit translation initiation has recently been identified in mammalian cells, although this specific mechanism is not thought to operate in yeast (Hickey et al. 2020; Juszkiewicz et al. 2020; Sinha et al. 2020). Recent work has demonstrated that slow elongation of non-optimal reporters can limit initiation to decrease protein output in human and fly cells (Barrington et al. 2023). Growing evidence suggests that slow decoding of natural transcripts can feed back to limit both the initiation and elongation stages of translation (Chu et al. 2014, Lyu et al. 2021).

Here, we investigate how elongation is rate limiting in protein production in yeast across the full range of codon optimality seen in native yeast genes. We use reporter assays and a high-throughput CRISPR interference screen to elucidate the scenarios linking slow elongation to lower protein output. Our results show that the consequences of slow elongation depend substantially on the rate of translation initiation, with a tipping point between these two potentially rate limiting processes. We establish that within the range of initiation and elongation rates of endogenous genes, protein output per mRNA is often limited by translation elongation, counter to long-standing models.

## Results

### Non-optimal reporters produce less protein per transcript, beyond the effect of mRNA decay

We first confirmed the large difference in observed fluorescence between our synonymous reporters. Using the six synonymous sequences of the yellow fluorescent protein citrine from our earlier work, we constructed yeast strains each containing one citrine reporter variant and a standard mCherry normalizing reporter. The reporters all employed the same promoter and 5′ untranslated region from the PGK1 gene, differing only in their coding sequence. The coding sequences were chosen based on their predicted total elongation time, calculated from a model of ribosome dwell time based on the mRNA sequence in and near the A site of the ribosome (Fig. 1A). The model designed the fastest and slowest possible encodings of citrine, and four sequences were chosen to represent the quartiles of a set of 100,000 randomly designed hypothetical sequences. The five fastest sequences — all but the overall slowest sequence — spanned the full range of predicted translation elongation rates of natural yeast genes (Fig. 1B). We confirmed that the protein output of the synonymous reporters closely tracked their predicted elongation time, with the fastest reporter producing 12 times more fluorescence than the slowest (Fig. 1C).

**Figure 1:**
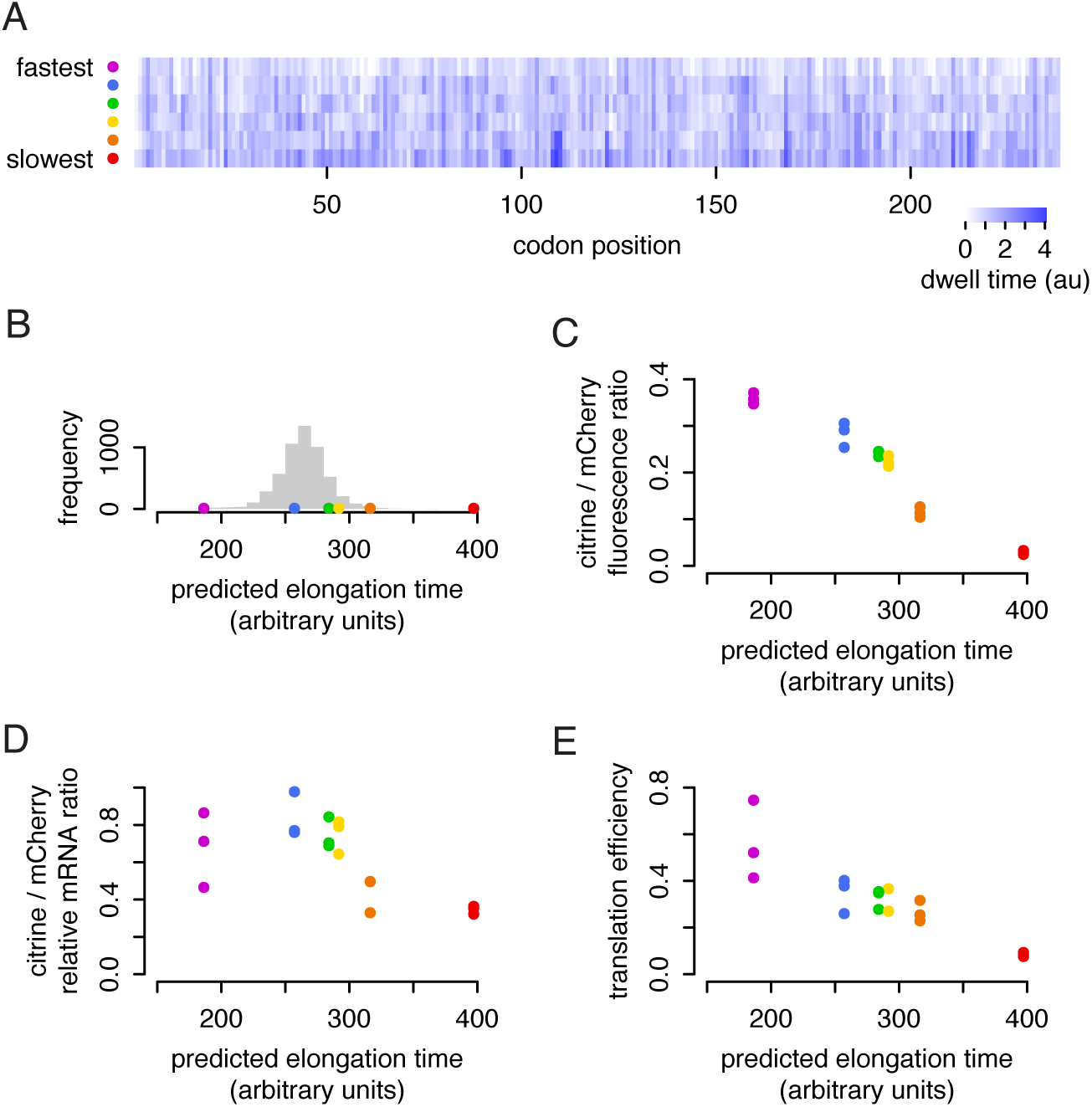
Non-optimal reporters produce less protein per transcript, beyond the effect of mRNA decay. a) Six synonymous citrine reporters depicted with their predicted ribosome dwell times at each position. Reporter sequences were designed in our previous work (Tunney et al, 2018), using a neural network model that predicts the time the ribosome spends on each position of a coding sequence as a function of the identity of the codon being decoded in the ribosomal A site and the adjacent codons in the P and E sites and neighboring sequence. The reporters were designed to span the range of total decoding times from fastest to slowest possible. b) The distribution of total predicted decoding times of endogenous yeast genes, normalized by length, is shown in gray, with the total elongation times of the six citrine reporters shown by color. c) Flow cytometry measurements of citrine fluorescence normalized to mCherry fluorescence for each synonymous reporter. Median fluorescence ratio for ∼20,000 yeast is shown for three isolates of each reporter; error bars per sample are too small to plot. d) mRNA abundance of each synonymous citrine reporter normalized to mCherry mRNA abundance, determined by RT-qPCR, for three isolates of each reporter. e) Translation efficiency of each synonymous reporter, calculated as the normalized fluorescence divided by normalized mRNA abundance.

To see if the lower output was due to destabilization of non-optimal transcripts, we then measured the mRNA abundance of each citrine reporter with RT-qPCR. When normalized to mCherry mRNA, citrine mRNA abundance also closely tracked the predicted elongation time, but only across a narrower 2-fold range (Fig. 1D). Non-optimal reporter transcripts were indeed less abundant, in keeping with the known relationship between slow codons and faster mRNA turnover (Presnyak et al. 2015), but mRNA abundance differences did not drive the majority of the difference in protein output. Dividing the total protein by total mRNA, we found that the translation efficiency, or proteins produced per mRNA in a given amount of time, spanned a 6-fold range (Fig. 1E). Thus, additional effects beyond mRNA decay had contributed to decrease the protein output from the non-optimal reporters.

To explore these effects, in much of this study we will focus specifically on the output of our fastest and second-slowest reporters, which we refer to as ‘fast’ and ‘slow’ citrine, respectively. These two reporters span the range of codon usage of endogenous genes, while the absolute slowest reporter is well outside of this range (Tunney et al. 2018). In the experiments above, the ‘fast’ reporter had twice the translation efficiency of the ‘slow’ reporter: each transcript generated twice as much functional mature protein per time (Fig. 1E).

### In a simple model, optimal and non-optimal reporters would have similar translation efficiency

To confirm our understanding that differences in elongation speed on their own should not produce different amounts of protein, we implemented a stochastic TASEP (Totally Asymmetric Simple Exclusion Process) simulation (Fig. S1A) (MacDonald and Gibbs 1969; Zia et al. 2011). Ribosome initiation rates and per-codon elongation rates were chosen to span a wide range of realistic values gathered from experimental data (Arava et al. 2003; Ingolia et al. 2011; Wang et al. 2022), and we simulated a shorter half-life for the slow reporter to reflect the impact of codon-optimality mediated decay and match its observed mRNA abundance. In this simulation, the translation efficiency for the slow reporter was predicted to be 92-93% that of the fast reporter (Fig. 2A; Fig. S1B). This stood in contrast to our experimental results, in which the translation efficiency of the slow reporter was only half that of the fastest reporter.

**Figure 2:**
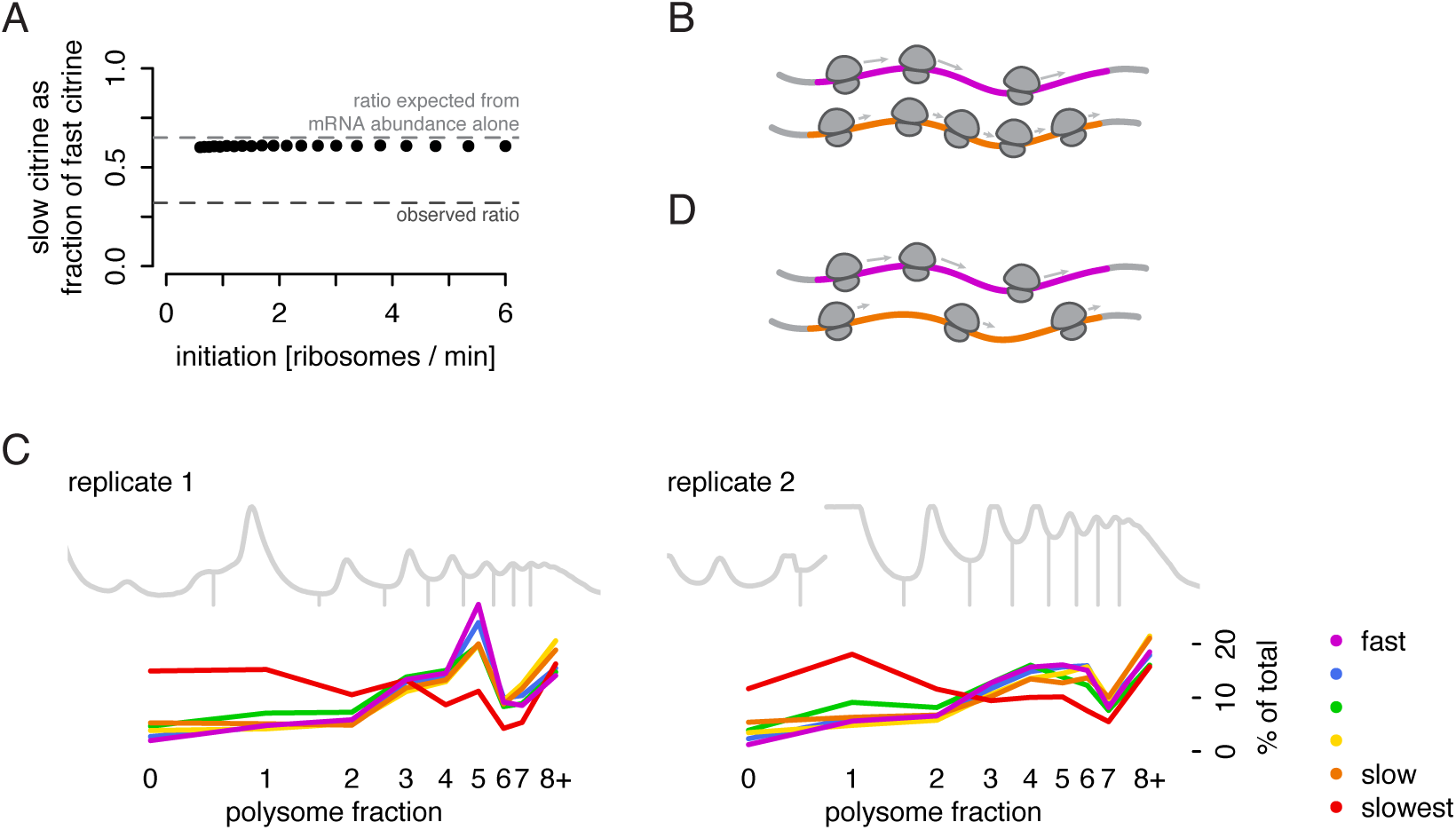
Measured translation output does not match predictions of a simple simulation. a) Simulated protein output from the TASEP model, shown as the ratio of slow reporter to fast reporter protein output, does not approach the observed ratio of protein output (dark gray dashed line) across a range of realistic initiation rates. Also shown is the protein output expected based only on mRNA differences (light gray dashed line). b) Schematic representation of expected ribosome density on fast (magenta) and slow (orange) synonymous reporter transcripts if no additional regulation beyond mRNA decay occurs to limit translation of the slow reporter. c) Distribution of ribosome counts on reporter mRNAs as determined by sucrose gradient fractionation of polysomes and RNA sequencing. To accurately compare reporter abundances in each fraction, a constant mass of human RNA was spiked into each fraction before RNA isolation. Abundance of each reporter in each fraction of the gradient is presented as a percentage of the total mRNA for that reporter. All six reporter strains were co-cultured to allow measurement from a single gradient; two replicates were performed with distinct isolates of the reporter strains. Polysome trace is depicted as 254 nm absorbance, gray. d) Alternative model explaining observed ribosome density on fast (magenta) and slow (orange) synonymous reporter transcripts.

This simple model has another implication: equal initiation, but slower elongation, should lead to accumulation of ribosomes on a transcript without affecting the total protein production per time. If an equal number of ribosomes initiate, but they take longer to reach the end of the coding region on a transcript with slow codons than on one with fast codons, then a slow citrine mRNA would generally be associated with more elongating ribosomes than a fast citrine mRNA (Fig. 2B). The total elongation time of our slow citrine reporter is predicted to be 70% longer than that of our fast citrine reporter, and we would expect its mRNAs to be occupied by commensurately more ribosomes. The TASEP simulations confirmed that, in this simple model, a slow citrine mRNA should be occupied by 50-60% more ribosomes than a fast citrine mRNA, with a notable difference in the mode of the distribution.

### Lower than expected ribosome density on non-optimal transcripts implicates a mechanism limiting translation

Counter to the simple models described above, our previous experiments showed that protein output is indeed limited from our slow reporters. To test whether the distribution of ribosomes also differed from the simple model, we went on to directly measure the ribosome density on our reporters with polysome profiling. We co-cultured yeast expressing all six reporters, used ultracentrifugation through a sucrose gradient to separate mRNAs by the number of associated ribosomes, then measured the abundance of each citrine mRNA in each polysome fraction via RNA sequencing. The results were striking: the five fastest constructs, ranging from our fast to slow citrine, all had a characteristic number of around 5 ribosomes per mRNA, with no shift in the mode, counter to predictions from simulations (Fig. 2C). Two small trends did depend on translation speed: the slower the sequence, the flatter the distribution, with a slight increase in mRNAs with no ribosomes and a slight increase in mRNAs with eight or more more ribosomes (these heavy polysomes are pooled into one fraction by necessity, truncating the distribution and appearing as a spike in the plot). In all, the number of translating ribosomes on the slower reporters was notably lower than the expected number (Fig. 2D). In contrast to the results on these five reporters, which span the endogenous range of codon usage, our slowest reporter showed severely limited translation, with many more mRNAs associated with just one ribosome or no ribosomes. The overall codon usage and speed of translation elongation on this reporter were far slower than that seen in any endogenous gene, suggesting that its sequence may trigger quality control mechanisms for defective transcripts that do not affect the other reporters that are in the endogenous range. The polysome distribution of the five faster constructs points to some mechanism limiting translation of normal but non-optimal mRNAs, and it is this unexpected impact of ‘normal’ slow elongation that interests us.

Our results thus led us to consider how slow translation elongation could determine protein output per mRNA. They point to some mechanism reducing initiation on slow transcripts, removing ribosomes from slow transcripts (beyond the effects of codon optimality mediated decay), or destabilizing the final protein product.

### Known quality control pathways do not explain decreased protein output

Aberrant translation has been shown to trigger surveillance pathways that can result in incomplete translation, and we reasoned that these pathways could also act on transcripts that are decoded slowly but without serious aberrations. To test if known translational quality control pathways were active on our reporters, we deleted a number of key genes in yeast that expressed either the fastest (‘fast’) or second-slowest (‘slow’) citrine reporter along with a normalizing mCherry reporter. We deleted genes encoding factors that are known to associate with or act downstream of ribosome collisions, including the no-go / non-stop decay ribosome collision detectors *HEL2* and *MBF1* (Sitron et al. 2017; Matsuo et al. 2017; Sinha et al. 2020), integrated stress response ribosome collision sensor *GCN1* (Sattlegger and Hinnebusch 2005; Pochopien et al. 2021), and E3 ubiquitin ligase *RKR1* (mammalian *LTN1*) which tags nascent chains for degradation in ribosome quality control (Bengtson and Joazeiro 2010). We also deleted *SYH1* and *SMY2* and made double knockouts of *SYH1* with both *SMY2* and *HEL2* to confirm the absence of regulation homologous to the mammalian GIGYF2-4EHP pathway in which collisions limit translation initiation (Hickey et al. 2020; Veltri et al. 2022). We observed no difference in normalized citrine fluorescence in any knockout strains compared to the original citrine strains (Fig. 3A), indicating that these quality control pathways are not actively reducing the protein output of our non-optimal reporters. The results suggest that our synonymous reporters, designed to capture the normal endogenous range of codon usage, are not subject to quality control mechanisms that respond to aberrant translation. Rather, an as-yet-uncharacterized mechanism may limit the number of ribosomes completing protein synthesis, either through repression of initiation or unsuccessful translation.

**Figure 3:**
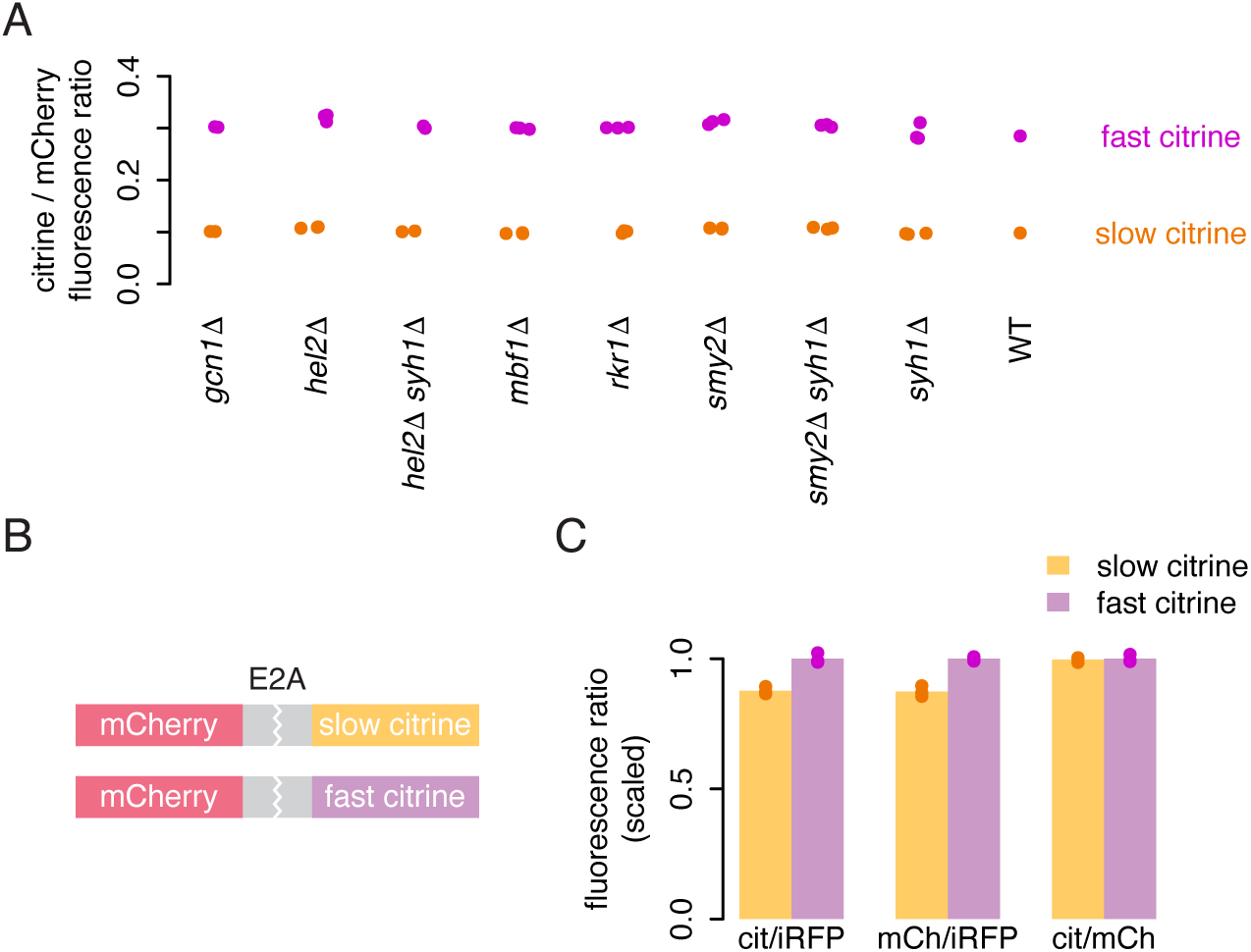
Lack of evidence for incomplete translation on non-optimal sequences. a) Flow cytometry measurements of fluorescence from fast and slow citrine reporters, as in figure 1B, in strains with deletions of genes implicated in ribosome associated quality control and related mechanisms. b) Schematic of dual fluorescence E2A constructs composed of an mCherry coding sequence with no stop codon, an E2A skipping sequence, and either the slow or fast citrine sequence. Yeast backgrounds also contain emiRFP for normalization. c) Flow cytometry measurements of citrine and mCherry fluorescence from fast and slow E2A constructs, each normalized to iRFP, as well as the ratio of citrine to mCherry. Citrine and mCherry measurements were each scaled to the average citrine or mCherry fluorescence, respectively, of the fast citrine reporter. The bar height shows the average of three isolates.

### Incomplete translation does not explain lower output of non-optimal sequences

To differentiate between these possibilities, we designed dual fluorescent reporter constructs with an upstream mCherry sequence followed by a viral E2A skipping element and either our fast or slow citrine variant (Fig. 3B). The E2A element causes ribosomes to skip catalysis of one peptide bond, releasing the upstream product while translation proceeds into the second sequence (Donnelly et al. 2001). Mechanisms by which slow elongation limits translation initiation or destabilizes the mRNA would reduce mCherry output to the same extent as citrine output from these E2A constructs. If, on the other hand, slow elongation caused some uncharacterized effect such as ribosome removal or destabilization of citrine protein, the effect would be decoupled, with less effect on mCherry output than citrine output.

We expressed the two E2A constructs in yeast along with iRFP as a normalizer and measured fluorescence via flow cytometry (Fig. 3C). As expected, the slow citrine E2A construct produced less citrine fluorescence than the fast citrine E2A construct, although the difference was attenuated relative to our original reporters. (This could indicate that the skipping sequence itself creates a translation stall and collision, decreasing output of both the slow and fast reporters (Wu et al. 2020).) Importantly, we observed that the slow citrine construct also produced less mCherry fluorescence, and that the ratio of citrine to mCherry fluorescence for both the fast and slow E2A constructs was identical. This indicates that the reduction in output caused by slow ribosomes on the downstream sequence is imposed equally on both the upstream and downstream sequences. This result is not consistent with the possibility of ribosome removal from the non-optimal citrine sequence or a difference in stability of the protein products from fast vs slow citrine sequences. Instead, it provides further support for a combination of mRNA destabilization and impaired initiation limiting output from slowly translated sequences. In the absence of ribosome removal, there must be fewer ribosomes initiating on slow reporter transcripts to generate the similar polysome profiles of fast and slow reporter mRNAs. Given these results, we considered mechanisms that might limit initiation on our non-optimal reporters.

### Passive start codon occlusion does not explain lower output of non-optimal sequences

In some cases, translation initiation can be limited by physical occlusion of the start codon by a ‘traffic jam’ of slowly elongating ribosomes (Plotkin and Kudla 2011; Shah et al. 2013; Chu et al. 2014; Tuller et al. 2010; Riba et al. 2019; Verma et al. 2019). A ribosome covers a footprint of around 30 nucleotides; any ribosome decoding the first ten codons of a transcript would block access to the start codon and interfere directly with initiation (Fig. 4A). This simple passive mechanism, in which slowly elongating ribosomes directly interfere with initiation, could potentially explain the lower protein output of our non-optimal reporters.

**Figure 4:**
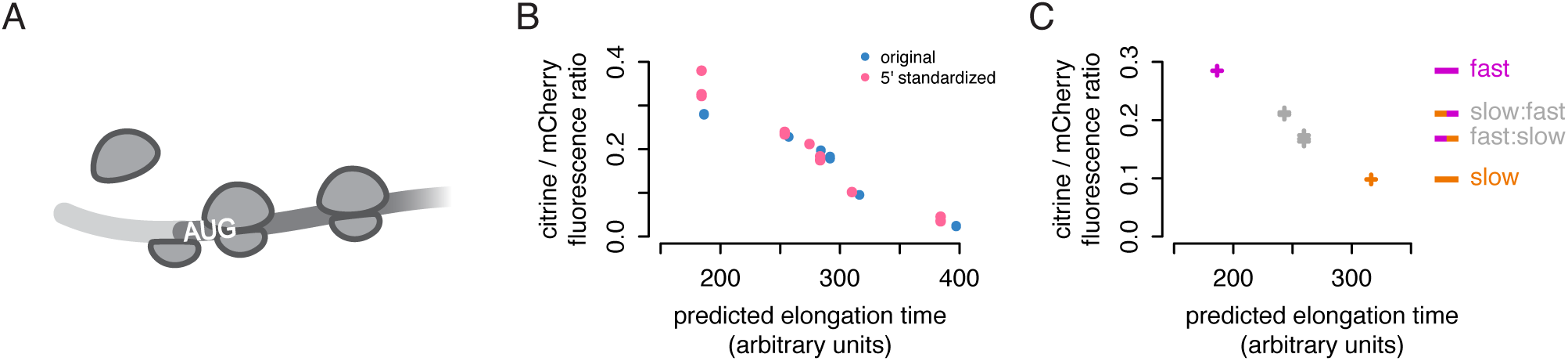
Passive start codon occlusion does not explain lower output of non-optimal sequences. a) Schematic of an initiation interference event; an upstream ribosome cannot initiate because the start codon is occluded by a downstream ribosome. b) Flow cytometry measurements of fluorescence from 5′-standardized and original synonymous citrine reporters, as in figure 1B. The first 20 codons of each reporter were standardized to the original yeCitrine sequence. c) Flow cytometry measurements of fluorescence from chimeric, fast, and slow synonymous reporters, as in figure 1B, from three isolates of each chimera and one of each original reporter. The reporters comprise the first half of the slow citrine sequence and the second half of the fast citrine sequence and vice versa.

Although ‘traffic jams’ are often proposed as an explanation of rate-limiting elongation, our TASEP simulation suggested this was unlikely to explain the differences in output between our reporters. Simulations predicted that the start codons on the slow reporter would be occluded only marginally more often than start codons on the fast reporter, with at most a 1% reduction in protein from the slow reporter when initiation was modeled to be particularly fast (Fig. S1C). This is clearly inconsistent with the 50% reduction in translation efficiency of the slow reporter relative to the fast reporter in our experiments. The TASEP results indicated that passive traffic jams were unlikely to explain the effect of slow codons on protein output.

To test experimentally whether the difference in output could be explained by the choice of codons proximal to the start codon, we created new versions of each of our six synonymous reporters in which the first 20 codons were standardized to the yECitrine sequence. We scored these new versions with our neural network model, which does not consider the position of a codon in a transcript; it essentially sums the decoding times of all positions in the transcript, with consideration of the impact of adjacent codons. Thus, its prediction of the total protein output of each 5′-standardized reporter only increased by a small percent relative to the original reporter. If codons near the start codon played a central role in limiting initiation, the experimentally measured output would diverge substantially from the neural network prediction; the 5′-standardized sequence would cause fewer traffic jams on the slower reporters and a concomitant increase in protein output. Instead, we observed only a small increase in the ratio of citrine to mCherry fluorescence for all synonymous reporter variants (Fig. 4B). The increase tracked with the slight improvement in predicted elongation time; the 5′-standardized reporters fit the previous linear trend between total predicted elongation time and protein output. This demonstrates that the protein output of our synonymous reporters is not determined by their 5′ sequence and argues against a 5′ ribosome traffic jam interfering with initiation and limiting output.

### Position of non-optimal codons does not influence protein output

We next asked more generally whether the position of non-optimal codons affected protein output. We recombined opposite halves of our fast and slow citrine reporters to generate slow:fast and fast:slow chimeric variants. We reasoned that if non-optimal codon position were relevant, the chimeric reporters would not track the pre-existing trend between predicted elongation time and protein output, as the elongation time prediction does not take codon position into account. Instead, we observe that the ratio of citrine to mCherry fluorescence closely tracked each chimeric citrine variant’s predicted elongation time (Fig. 4C). The 5′-non-optimal slow:fast variant displayed slightly higher output than the 5′-optimal fast:slow variant, due to an overall higher fraction of fast codons in the former. This result indicates that there is no strong positional effect of non-optimal codons within these coding sequences.

Taken together, these findings suggest that physical occlusion of the start codon by ribosome traffic jams is not involved in reducing protein output from our non-optimally encoded synonymous reporters. We have established clearly that our slower reporters produce less protein per mRNA and are occupied by fewer ribosomes than expected by a simple model. With evidence against incomplete translation and start-codon-occluding traffic jams, we considered the most plausible remaining explanation to be additional, uncharacterized mechanisms that reduce translation initiation on non-optimal transcripts.

### A CRISPRi screen identifies initiation factors as modulators of non-optimal codon translation

To identify factors that determine whether slow codons limit protein output, we screened for genes whose inhibition caused differential effects on our fast versus slow citrine variants. We modified a pooled, genome-wide CRISPR interference screen, CiBER-seq (Muller et al. 2020), by fusing a citrine variant sequence to a synthetic transcription factor that drove expression of a transcriptional reporter (Fig. 5A; Fig. S2A). Changes in translation of the citrine sequence would change the abundance of the citrine-transcription factor fusion and in turn change the expression of the reporter transcript. A pool of plasmids, each containing a transcriptional reporter with a unique barcode and a Cas9 guide RNA against the promoter region of one yeast gene, were transformed into a population of yeast, allowing unique identification of the CRISPRi target in the cell expressing a particular barcode. By sequencing the reporter mRNAs across all cells and counting the frequency of each barcode, we can measure how knockdown of each gene influences citrine translation with greater sensitivity and precision than a standard approach of sorting yeast by fluorescence output.

**Figure 5:**
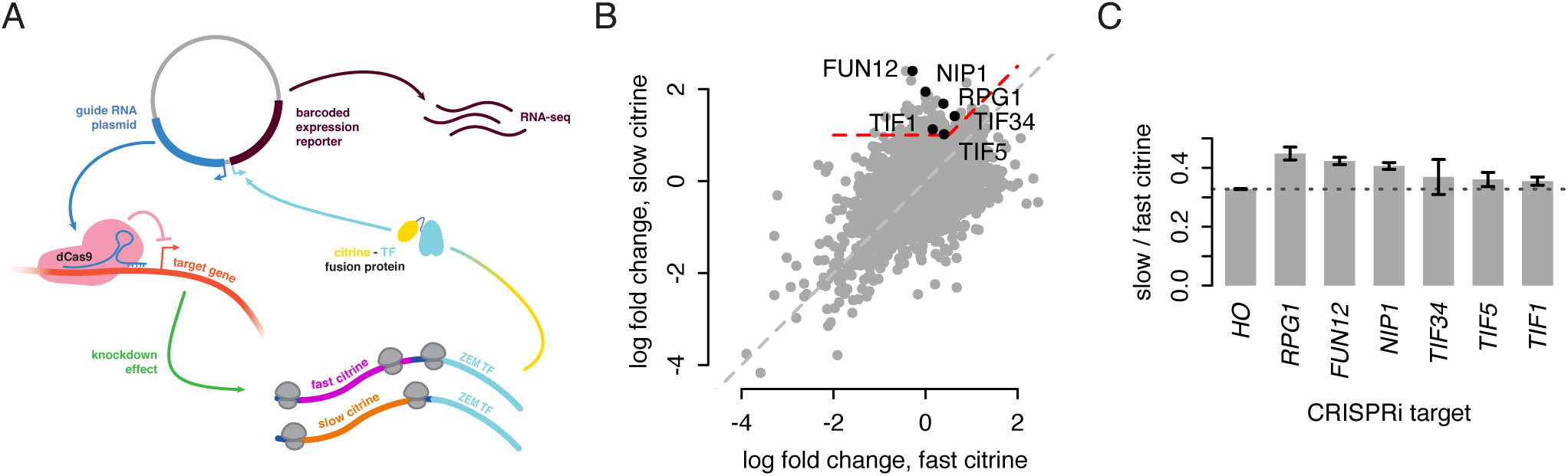
A CRISPRi screen identifies initiation factors as modulators of non-optimal codon translation. a) The modified CiBER-seq assay. Each Cas9 guide RNA cassette is linked to a transcribed reporter with a unique barcode sequence on the same plasmid. Expression of this barcoded reporter is driven by the synthetic transcription factor ZIF286 (ZEM-TF) fused downstream of either the fast or slow synonymous citrine sequence. The RNA-to-DNA ratio for each barcode, determined by deep sequencing, corresponds to reporter expression driven by the citrine-ZEM fusion. A change in translation of the citrine sequence due to CRISPRi knockdown of a relevant gene will result in changes in citrine-ZEM abundance and expression of the barcode linked to that guide RNA. b) Log_2_ fold change (LFC) of barcode counts after guide induction for fast citrine (x-axis) and slow citrine (y-axis) fusion backgrounds, for each linked guide. Red dashed line indicates the region containing guides with linked barcodes that showed LFC > 1 in the slow citrine fusion background and (LFC_slow_ - LFC_fast_) > 0.5. Labels indicate guides that correspond to the term ‘cytoplasmic translation initiation,’ the only overrepresented term in a Gene Ontology analysis of all guides in the region of interest. c) Ratio of normalized fluorescence of the slow citrine reporter to normalized fluorescence of the fast citrine reporter in isogenic CRISPRi backgrounds. Dashed line indicates the value of this ratio for the HO guide background, which is used as a control. Fluorescence was determined by flow cytometry; bar height corresponds to the average of three isolates of the citrine reporter in each guide background and error bars depict standard error of the mean.

To find differential effects on fast and slow citrine translation, we transformed the plasmid library into yeast containing either the fast or slow citrine-transcription factor fusion and compared outcomes (Fig. 5B). To identify genes whose inhibition allowed for higher relative expression of slow citrine without a substantial effect on fast citrine, we analyzed the 118 guides with a two-fold increase in reporter count in the slow citrine strain and a large difference between the fast and slow citrine strains (Δ(log2 fold change) > 0.5). We reasoned that genes displaying this effect upon knockdown would contribute to translational repression of non-optimally encoded sequences. Notably, no guides against known translational quality control genes showed a large difference in effect between the fast and slow citrine backgrounds. Instead, the data revealed an impact of translation initiation. We found two significantly overrepresented Gene Ontology terms within this set: ‘cytoplasmic translation initiation’ (p = 0.03, hypergeometric test with Benjamini-Hochberg false discovery rate < 0.05) and the more specific ‘formation of cytoplasmic translation initiation complex’ (p = 0.002). The first term was represented by six initiation factor genes: *RPG1*, *NIP1*, and *TIF34* (eIF3 subunits a, c, and i); *FUN12* (eIF5B); *TIF1* (eIF4A); and *TIF5* (eIF5). The second term included all these genes except eIF4A. While various steps of translation initiation were implicated, the presence of three subunits of eIF3 could suggest a specific role for this complex, and we note that a guide against the g subunit *TIF35* also showed a strong differential effect below our threshold.

### Initiation factor depletion reduces difference in synonymous reporter output

We verified the results of the CiBER-seq screen by measuring the effect of initiation factor knockdown on citrine translation more directly using flow cytometry. We created individual yeast strains with either the fast or slow citrine reporter, CRISPRi machinery, and inducible guides against each of the six initiation factors and against the HO locus as a control for general effects of CRISPRi. To confirm that our results were not a general result of growth defects, we also tested two guides that would cause large growth defects but showed no differential effect in the CiBER-seq screen (Fig. S2B). We compared the ratio of slow to fast citrine fluorescence between the initiation factor targets and the HO control and found that knocking down all targets except *TIF34* showed an increase in the fluorescence ratio, indicating a decrease in the difference between slow and fast output (Fig. 5C). We confirmed that knockdown of *RPG1* affected translation, not mRNA stability, as reporter mRNA levels did not change (Fig. S2C). We conclude that, while translation of all genes presumably decreases when initiation factors are depleted, this decrease is less pronounced for our slow reporter. Thus, the impact of non-optimal codons on translation is lessened when initiation is disrupted.

Translation initiation was clearly implicated, and our next goal was to distinguish between two possibilities. Our knockdowns could have disrupted a specific active mechanism that responds to non-optimal codon translation by altering translation initiation. But, the results also suggested a more general possibility: lower impact of slow codons could be a direct consequence of having fewer translating ribosomes after initiation factor depletion. In other words, the potential for slow codons to limit the rate of protein production could itself depend on the rate of initiation. For instance, ribosome collisions, the activating event for many quality control pathways, would not occur as frequently on transcripts with lower initiation and fewer translating ribosomes. Our reporters were constructed with the 5′ UTR of PGK1, a highly expressed gene with high translation efficiency, but the initiation rate varies widely on endogenous transcripts, driven in part by differences in 5′ UTR sequence (Niederer et al. 2022, Akirtava et al. 2024). The impact of slow codons in endogenous transcripts could be different in the context of lower initiation. This led us to wonder if disrupting initiation in an orthogonal manner would result in a similar change in the relative translation efficiency of our synonymous reporters.

### Lowering initiation with 5′ UTR stem-loops attenuates the difference in protein output

To probe how translation initiation rate affects the rate-limiting step in protein output from our reporters, we inserted RNA stem-loop (SL) structures with different folding energies, strong-SL and weak-SL, directly upstream of the start codon in each citrine construct (Fig. 6A). These stem-loop structures have been shown to reduce protein expression in proportion to their folding energy (Weenink et al. 2018).

**Figure 6:**
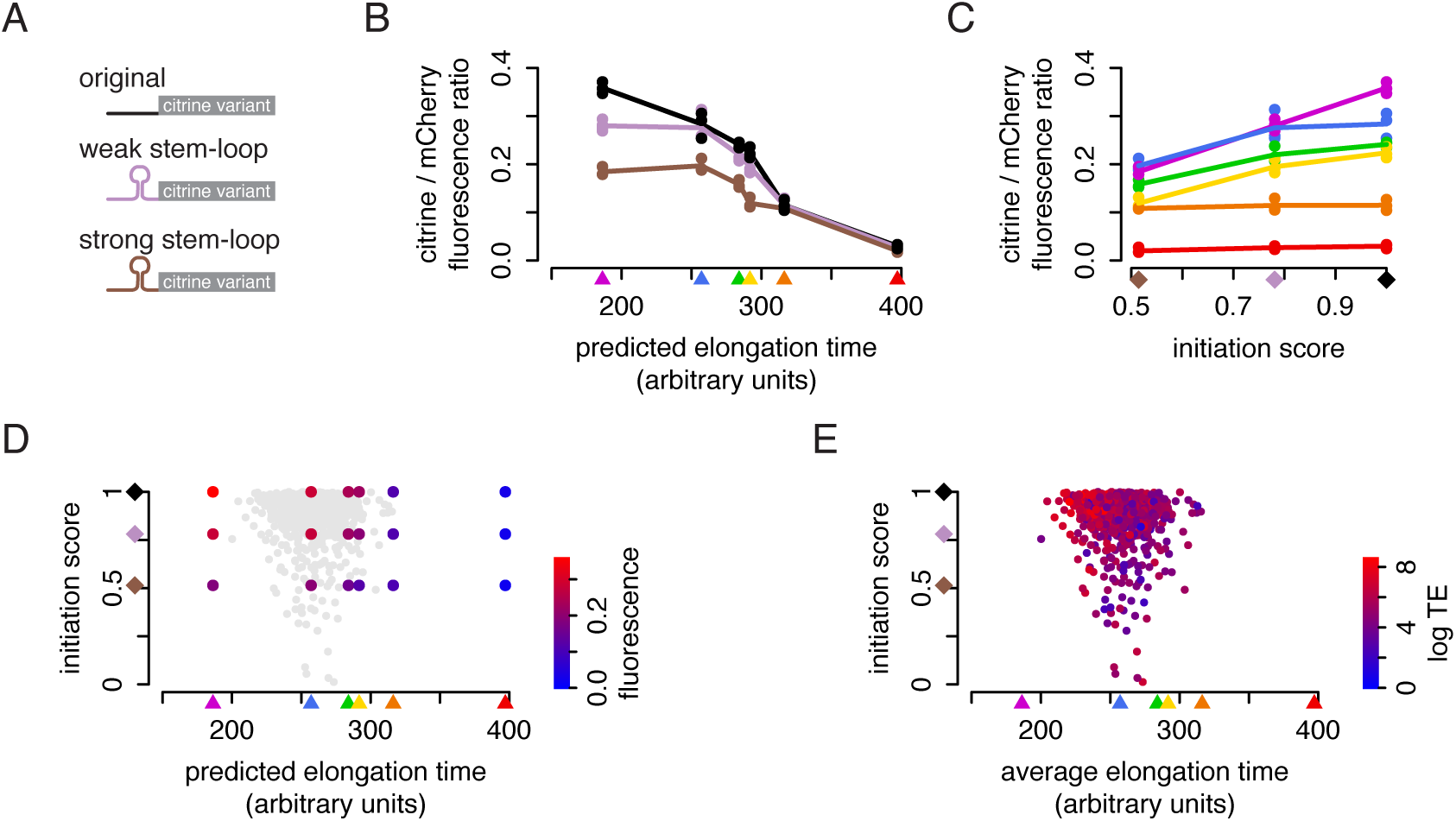
The impact of non-optimal codons on protein output depends on initiation rate. a) Synonymous citrine reporters were combined with 5′ UTRs with and without stem-loop structures, creating 18 reporter combinations. b) Flow cytometry measurements of fluorescence from the reporters. Points represent the median yellow:red ratio of ∼20,000 cells. Lines represent the average of three replicates. c) Protein output from the fastest reporter was used to compute an initiation score for each UTR variant as its output relative to the variant with no stem-loop. Fluorescence output of the six citrine coding variants, calculated as in (b), is plotted as a function of this initiation score. Lines represent the average of three replicates. d) Fluorescence output of the 18 reporter variants, represented by a color gradient, as a function of initiation score and predicted elongation time. Value represents the average of three replicates. Gray dots show the genes analyzed in (e). e) A set of ∼1000 endogenous yeast genes were plotted by the average elongation time of each endogenous CDS, as predicted by our iXnos method, and by the output driven by each endogenous UTR when fused to a YFP reporter (Akirtava et al, 2024). Points were colored by the translation efficiency (proteins per mRNA per time) calculated from measurements of protein abundance, protein turnover, and mRNA abundance (Lahtvee et al, 2017).

We measured the fluorescence of the stem-loop reporter variants with flow cytometry, normalizing to the mCherry reporter (Fig. 6B). We observed that the stem-loops had a varied, non-linear effect across our reporters. The weak stem-loop lowered the output of only the fastest reporter, while the strong stem-loop lowered the output of all but the two slowest reporters. Overall, we saw a remarkably clear shift from initiation-limited to elongation-limited translation as elongation speed became slower or initiation became faster.

We sought to quantify the transition between these two regimes of rate-limiting steps of proteins synthesis by considering the output of each citrine more explicitly as a function of its initiation rate. We computed initiation scores for the three UTRs based on their output when fused to the fastest citrine as a fraction of the maximum. If output of the fastest citrine was not limited at all by its codon content, this score should approximate the intrinsic initiation rate of each UTR. Plotting the output of each citrine variant as a function of this initiation score emphasized that the two slowest citrines were entirely limited by elongation, with no impact of initiation; output of the fastest citrine was very sensitive to changes in initiation; and output of the citrine reporters with intermediate codon usage, in the range of many endogenous genes, depended on both contributions (Fig. 6C). Considering the fluorescence output of each of the reporters as a function of both elongation and initiation defines the space of possible contributions of codon choice and UTR design on protein output, with areas of this space limited more by one or the other aspect (Fig. 6D).

We next considered how our measurements aligned with the output of endogenous genes. We obtained careful estimates of protein output per mRNA per time for ∼1000 yeast metabolic enzymes from a study which measured protein level, mRNA level, and protein turnover rate (Lahtvee et al. 2017). We predicted the average elongation time for the coding region of each gene with our iXnos model, and we obtained initiation scores for the endogenous 5′ UTR of each gene from a study that measured output of each endogenous UTR fused to a yeast-optimized yellow fluorescent protein (Akirtava et al. 2024). The protein produced per mRNA per time showed a correlation with the elongation time of its CDS (Spearman’s ρ = −0.35) as well as with the initiation score of its UTR (Spearman’s ρ = 0.17) (Fig. 6E), suggesting that across the physiologic range of combinations of initiation and elongation rates, protein output per mRNA is frequently limited by translation elongation.

We find that the median average translation elongation time of endogenous genes is similar to our second-fastest reporter (indicated in blue in Fig. 6). Our stem-loop reporter results predict that higher initiation would lead to higher output of these median genes up to some point, after which output would plateau. Many of these endogenous genes have UTRs with quite high intrinsic initiation rates (Fig. 6E), and this would suggest that they are in a regime where they are not getting the full benefit of their initiation rate and that instead, codon choice is limiting their output. In support of this, our analysis of endogenous genes shows that the translation efficiency of genes with faster-than-median elongation correlates slightly more strongly with the UTR strength (Spearman’s rho = 0.21), while for genes with slower-than-median elongation, the correlation is lower (Spearman’s rho = 0.15). Our carefully designed combinations of synthetic reporter sequences thus reveal constraints on the translation of natural genes, establishing that both initiation and elongation can limit protein output through unappreciated mechanisms.

## Discussion

Paradoxically, the speed of ribosomes moving along an mRNA should not necessarily affect the number of proteins made per time. Unless ribosomes move slowly enough to cause a traffic jam and block initiation of more ribosomes, the intrinsic initiation rate should determine eventual protein output. In the absence of feedback mechanisms, the number of ribosomes initiating and the number completing translation should match. mRNA decay is known to be enhanced by slow elongation, limiting output from non-optimal transcripts, but each surviving mRNA should still produce approximately the same amount of protein per time.

Nonetheless, in this study we have established that codon choice and translation elongation speed can directly determine translation efficiency and protein output in yeast, across a wide range of codon optimality. Using a series of synonymous fluorescent reporters designed to span the full range of elongation speeds, we found that their protein output scales linearly with total elongation time. Our polysome analysis shows conclusively that mRNAs with slow codons are translated by fewer ribosomes than would be expected without additional feedback mechanisms. Importantly, this effect is not primarily explained by the known impact of slow decoding on mRNA turnover. Each slow mRNA produced fewer proteins than a fast mRNA; some mechanism must limit protein output when translation elongation is slow.

We have systematically investigated the ways that the speed of elongation might limit protein output from an mRNA: by causing fewer ribosomes to start translation, causing fewer ribosomes to successfully complete translation, or creating defective or unstable proteins. Unlike transcripts with severe translation stalls or other defects (Veltri et al. 2022; Hou et al. 2023), we found that our slow reporter did not trigger quality control pathways. (Our experiments intentionally focused on the reporters that span the range of endogenous codon usage, excluding the overall slowest reporter that showed notably different polysome association and may have severe translation defects.) We also ruled out an effect of passive ‘traffic jams’ occluding the start codon of the slower reporters and showed that the rate-limiting impact of slow codons on protein output does not depend on their location within a transcript.

The strongest remaining possibility is that some process limits initiation on mRNAs with slow codons. Very similar conclusions have recently been reached by Barrington et al. (2023) who found that translation was limited on non-optimal transcripts in human and fly, and showed that these transcripts were less associated with initiation factors including eIF4E and eIF4G1, indicating a mechanism limiting translation in response to slow elongation. The discovery of specific feedback on initiation in response to ribosome stalls (Hickey et al. 2020; Juszkiewicz et al. 2020; Sinha et al. 2020) makes it plausible that an analogous mechanism could function on slow but non-stalled ribosomes.

The results of our genome-wide CRISPRi screen emphasize the interplay between translation initiation and elongation. Translation initiation factors, including multiple subunits of eIF3, were enriched among the genes whose depletion reduced the difference in output between our fast and slow reporters. It is possible that these factors play a specific role in a feedback mechanism limiting initiation in response to slow elongation. Many studies have pointed to diverse roles for eIF3 across all stages of translation, including initiation (Martineau et al. 2008), elongation (Pöyry, Stoneley, and Willis 2020; Lin et al. 2020; Wagner et al. 2020; Bohlen et al. 2020) and termination (Beznosková et al. 2013), and its impact on initiation may go beyond its canonical role as a scaffold for pre-initiation complex formation. However, the results of our stem-loop reporter experiments point in a more general direction. Reducing initiation, whether globally or via UTR sequence, seems to attenuate the impact of slow elongation on protein output. This implies that the optimality of codons, and their ability to limit protein output, can only be interpreted in the context of the initiation rate they encounter, as has been proposed by Park and Subramaniam (2019).

To consider the rate limiting aspects of translation, we can ask what would happen if a particular gene were changed to have faster codons while keeping the same UTR sequence. If its protein output per mRNA increased, we could conclude that its codon choice was limiting output to some extent. The results from our reporters suggest that the outcome of this experiment would depend on the strength of the gene’s UTR. Genes with the strongest initiation would be sensitive to elongation speed across the range of codon choices seen in normal genes, while genes with weak initiation would not be affected much by changes in elongation speed. Conversely, at the fastest elongation speeds, initiation rate would have a large role in determining protein output, but at the slowest elongation speeds, initiation rate would matter very little. Many endogenous genes seem to fall in the middle of this range. It is interesting that the point at which initiation stops mattering as a function of elongation, and vice versa, both fall in the middle of the endogenous range of elongation and initiation rates, suggesting that this relationship has co-evolved (Park and Subramaniam 2019; Riba et al. 2019).

Global changes in translation initiation are a key part of many cellular pathways. Stress causes cells to phosphorylate key components of the translation initiation machinery and thereby decrease initiation of most genes, and similar processes are important for formation of long-term memories in neurons. A major implication of our results is that the proteome will respond non-uniformly to these global changes in initiation, with differential effects on production of proteins encoded with more optimal or less optimal codons. Rather than simply scaling protein production up or down, changing the global rate of initiation may reshape the proteome in unappreciated ways. The interplay between initiation and elongation shapes this regulatory program and has implications for the evolution of natural sequences and the design of synthetic mRNAs.

## Data availability

Analysis code and data are available at https://github.com/lareaulab/synonymous-citrines

## Acknowledgments

This work was supported by the National Institutes of Health (National Institute of General Medical Sciences R01GM132104 to L.F.L. and R01GM130996 to N.T.I.), the National Science Foundation (1936069 to L.F.L.), the Rose Hills Foundation (to L.F.L.), and the Chan Zuckerberg Biohub (to L.F.L.). M.V.M. was supported by a fellowship from the ARCS Foundation. We are thankful for the Vincent J. Coates Genomics Sequencing Laboratory (QB3 Genomics, UC Berkeley, Berkeley, CA, RRID:SCR_022170; supported by NIH S10 OD018174 Instrumentation Grant) for sequencing support, and for the U.C. Berkeley flow cytometry core facilities. We thank Joel McManus for sharing prepublication data and Robert Tunney, Meghan Pressimone, and Joe Lobel for their contributions.

## Declaration of interests

N.T.I. is a shareholder of Velia Therapeutics and a shareholder and member of the scientific advisory board of Tevard Biosciences. L.F.L. holds a patent on the iXnos mRNA design method used here.

## Methods

### Plasmid and yeast strain construction

The citrine variant sequences used and modified in this study were previously described in Tunney, et al. (2018). All fluorescent protein expression was directed by a *PGK1* promoter and an *ADH1* terminator.

Citrine variants were integrated into the yeast genome using the plasmids described in Tunney, et al. (2018), containing a *K. lactis LEU2* expression cassette and two 300bp sequences homologous to the *his3Δ1* locus of BY4742 for homologous recombination. pPGK1-mCherry was amplified from the mCherry plasmid described in Tunney et al. (2018) using uracil-containing primers with Q5U polymerase (New England BioLabs, M0515) and inserted into EasyClone plasmid pCfB2226 by USER cloning upstream of the *ADH1* terminator with USER Enzyme Mix (NEB, M5508) according to Stovicek et al. (2015). The resulting mCherry plasmid contains an *S. pombe HIS3* expression cassette for selection and integrates into the *S. cerevisiae* X-4 chromosomal locus.

All subsequent amplifications for cloning purposes used Q5 2X MasterMix (NEB, M0492). Knockout constructs were created by amplifying either a KanMX or HygR resistance marker from a parent plasmid with two sets of overlapping 60nt primer pairs, generating amplicons consisting of the selectable marker flanked by 80-100bp of target locus homology.

Dual fluorescence E2A reporter constructs were constructed as follows. The E2A sequence was divided into halves that overlapped by 15nt. Fast and slow citrine plasmids were linearized by amplification. The forward primer for each began at the citrine start codon and was tagged with the downstream half of the E2A sequence. The reverse primer began directly upstream of the start codon. Amplified plasmids were digested with DpnI (NEB, R0176). The mCherry coding sequence was amplified with a forward primer located at the mCherry start codon that was tagged with citrine vector homology. The reverse primer began one codon before the mCherry stop codon and was tagged with the upstream half of the E2A sequence. The mCherry insert was assembled into the citrine vector using NEB HiFi DNA assembly (NEB, E2621). The emiRFP reporter was constructed by linearizing the mCherry plasmid backbone with primers positioned to exclude the mCherry coding sequence. The emiRFP was amplified from a donor plasmid with primers containing mCherry backbone homology. The emiRFP insert was assembled into the mCherry backbone using NEB HiFi DNA assembly.

5′-standardized citrine variant plasmids were constructed by whole plasmid PCR of each variant using opposing primers containing the standardized codon sequence. Amplified plasmid was gel purified and then circularized with KLD Enzyme Mix (NEB, M0554).

Chimeric citrine reporter plasmids were constructed by amplifying each half of the fast and slow citrine sequences. A citrine plasmid was double digested at the PacI and AscI site to remove the citrine coding sequence from the backbone and treated with Quick CIP (NEB, M0525) to prevent recircularization. The up- and downstream halves of each chimeric citrine variant were assembled into the backbone using HiFi DNA Assembly MasterMix with a 60nt ssDNA oligo bridge according to manufacturer’s instructions.

Individual CRISPRi guide expression plasmids were constructed by inserting a tet-inducible guide expression cassette from Muller, et al. (2020) into EasyClone plasmid pCfB2188, containing the KanMX marker for selection, via USER cloning according to Stovicek et al. (2015). Individual guide sequences were ordered as 60nt ssDNA oligos containing the 20nt guide sequence flanked by 20nt backbone homology on either side, and were integrated into the AvrII site using HiFi DNA Assembly MasterMix.

Stem-loop citrine reporter plasmids were constructed by assembling four overlapping 60nt ssDNA oligos containing the stem-loop sequence into the PacI site of each citrine variant using HiFi DNA Assembly MasterMix according to manufacturer’s instructions.

All USER reaction products and HiFi assembly products were transformed into chemically competent *E. coli* cells (NEB 5-alpha: C2987, Agilent through UCB MacroLab XL1-Blue: 200249) according to the manufacturer’s directions. Plasmids were harvested from overnight cultures using either Zymo Miniprep Kits (Zymo, 11-308) or NEB Miniprep Kits (NEB, T1010).

All plasmids were sequence confirmed and transformed into yeast using the high-efficiency lithium acetate/single-stranded carrier DNA/PEG method according to Gietz and Woods (2002). Citrine plasmids were linearized at the SbfI site, and ∼1 µg linearized plasmid was used to transform BY4742 yeast. mCherry plasmid was linearized at the NotI site, and ∼1ug linearized plasmid was transformed into BY4741 yeast. Citrine transformants were selected by growth on SCD-Leu plates and mCherry transformants were selected on SCD-His plates. Knockout constructs were transformed into mCherry and citrine yeast backgrounds, which were then grown on YEPD plates overnight, then replica plated onto the appropriate antibiotic plates for selection. CRISPRi strains were constructed by transforming the dCas9-Mxi expression plasmid from Muller et al. (2020) into the mCherry yeast background and selected for on kanamycin plates. CRISPRi guide expression plasmids were linearized at the *NotI* site, and ∼1ug of linearized plasmid was transformed into the mCherry yeast background and selected for on SCD-Ura plates.

Transformant colonies were picked from selective plates and cultured overnight at 34°C in 1mL selective liquid media in deep-well 24-well blocks supplemented with one glass bead per well, shaking at 750rpm in a Multitron incubator shaker. Genomic integration of all constructs was confirmed by using 1uL of liquid culture in a colony PCR reaction with Phire Green Hot Start II PCR Master Mix (ThermoFisher, F126) with primers flanking the integration site. Genotyped transformant cultures were struck out onto selective plates and preserved as glycerol stocks in 30% glycerol at -80°C.

Diploid strains expressing both citrine and mCherry were created by inoculating 1mL of YEPD with one BY4741 strain and one BY4742 strain, which was then incubated overnight in 24-well blocks as described above.

Overnight mating cultures were then struck out onto SCD-Met-Lys plates for diploid selection. To create diploid knockout strains, three isolates of each haploid mCherry knockout strain were mated with separate isolates of each haploid citrine knockout strain, yielding a total of three diploid biological replicates with no parental isolate backgrounds in common. All other diploid yeast strains were created by mating one haploid isolate of mCherry yeast to three separate isolates of haploid citrine yeast, yielding three biological replicates that share the same parental mCherry isolate background.

### Yeast culture

Yeast strains were cultured overnight in either YEPD or selective dropout media, then back-diluted into YEPD to an OD_600_ of 0.1 and shaken at 250 rpm at 30°C in a sterile 24-well deep-well block supplemented with a sterile glass bead until their OD_600_ was between 0.4 and 0.7.

CRISPRi yeast strains were cultured overnight in SCD-Met-Lys, then back diluted into either SCD or YEPD to an OD_600_ of 0.1, and shaken at 250 rpm at 30°C in a sterile 24-well deep-well block supplemented with a sterile glass bead until their OD_600_ was between 0.4 and 0.7. Cultures were then serially back diluted into SCD or YEPD with tetracycline at final concentration of 250ng/uL, to an OD_600_ of 0.0005, 0.0001, and 0.00005, then grown overnight by the same method as above. Serial dilutions were necessary because CRISPRi strains have differing degrees of growth defect. CRISPRi-induced yeast were collected after at least 9 doublings at an OD_600_ between 0.4 and 0.7.

### Flow cytometry and analysis

200uL aliquots of suspended mid-log phase culture was pelleted by centrifugation at 3000*g* for 5 minutes in a 96-well U-bottom plate, washed with 200uL of DPBS (Gibco, 14190-44), fixed in 4% formaldehyde for 15 minutes in the dark at room temperature, then washed twice and diluted 1:4 in DPBS and stored at 4°C.

Flow cytometry of fixed yeast was carried out on a BD Biosciences LSR Fortessa analyzer using a High Throughput Sampler. Forward light-scatter measurements (FSC) for relative size and side-scatter measurements (SSC) for intracellular refractive index were made with a 488-nm laser. Citrine fluorescence was measured with 488-nm (blue) laser excitation and detected with a 505-nm long-pass optical filter followed by a 525/50 nm optical filter with a bandwidth of 50 nm. mCherry fluorescence was measured with a 561-nm (yellow–green) laser for excitation and a 600-nm long-pass optical filter followed by 610/20-nm band-pass optical filter with a bandwidth of 20 nm. emiRFP fluorescence was measured with a 561-nm (yellow–green) laser for excitation and a 635-nm long-pass optical filter followed by 670/30-nm band-pass optical filter with a bandwidth of 30 nm. PMT values for each color channel were adjusted such that the mean of a sample of BY4743 yeast was 100. 50,000 events were collected for each biological sample.

Flow cytometry data were analyzed with a custom R script (available on Github in “analysis code” folder) whose core functionality is based on the Bioconductor packages flowCore, flowStats, and flowViz. To select events that represented normal cells, we used the norm2filter method to extract events that had FSC and SSC values within the region of highest local density of all events. This gating method fits a 2D normal distribution and selects all events within the Mahalanobis distance, around 20,000 event per sample. For these gated events, the red and yellow fluorescence intensities were normalized by subtracting median red and yellow intensities from a control sample with no fluorescent reporters grown in the same media and conditions. For each sample, we computed the citrine to mCherry ratio as the median ratio of the background-corrected yellow and red intensities.

### CiBER-seq assay

#### Strain and plasmid construction and turbidostat culture

The inducible gRNA-barcode plasmid library, the citrine-ZIF286 fusion protein plasmid, and the dCas9-Mxi1-TetR plasmid used in the modified CiBER-seq assay were generated as described in Muller, et al. (2020). Yeast strains containing either a fast or slow citrine-ZIF286 fusion and the dCas9 plasmid were generated according to the yeast transformation protocol described above. These strains were then transformed separately with aliquots of the same pooled gRNA-barcode plasmid library, and serial dilutions were plated to ensure sufficient transformation efficiency was achieved. In all subsequent steps SCD-Ura media was used to maintain the non-integrating guide plasmid. Transformants were shaken overnight at 250 rpm in a 30° shaking water bath, then back diluted to an OD_600_ of 0.1, and allowed to grow until mid log phase. A custom made turbidostat (McGeachy, Meacham, and Ingolia 2019) was then inoculated with both fast and slow CiBER-seq yeast strains in two replicate growth chambers, and allowed to run continuously until an OD_600_ equal to 0.8 was reached. Media used in the turbidostat was supplemented with 8 nM beta-estradiol to allow ZIF286 nuclear import. When the growth rate reached steady state (90 minute doubling time), 50mL pre-induction samples were collected. gRNA expression was then induced by adding anhydrotetracycline in DMSO into both of the growth chambers and the media reservoir to a final concentration of 250 ng/ml. 50mL post-induction samples were collected six doublings later, after approximately 9 hours. Collected samples were centrifuged at 4000 × g for 5 min, the media was aspirated, and the pellets were stored at −80°C.

#### DNA and RNA library preparation

Plasmid DNA was extracted from yeast using Qiagen QIAprep Spin Miniprep Kits (Qiagen, 27104). 500uL Resuspension Buffer PI was added to thawed pellets, then 100µL acid-washed glass beads (Sigma, G-8772), followed by vortexing for 10 minutes. 500µL Lysis Buffer P2 was added, and tubes were gently inverted 4-6 times, then incubated at room temperature for 5 minutes. 700µL Neutralization Buffer N3 was added, and tubes were inverted 4-6 times, then centrifuged at maximum speed for 10 minutes. Cleared lysate was transferred to a spin column, flowed through, washed with 750µL Wash Buffer PE, and spun dry. DNA was eluted by adding 45uL pre-heated nuclease-free water (37°C), then incubating for 5 minutes at room temperature before centrifugation.

Eluate was digested with *PvuII-HF* (NEB, R3151S) for 2-3 hours, then column cleaned in a Zymo 5µg DNA Clean and Concentrate column (Zymo, D4004). In vitro transcription was performed using a T7 Quick HiScribe kit (NEB, E2050S) according to manufacturer’s instructions with overnight incubation at 37°C, then treated with DNase. IVT product was then reverse transcribed with a ProtoScript II kit (NEB, E6560S) according to manufacturer’s instructions, then treated with RNase A and RNase H. Purified product was amplified with distinct NEBNext dual index primer sets (NEB, E7600S) for 6 cycles with Q5 polymerase in 50uL reactions.

RNA was isolated from yeast using acid phenol chloroform extraction according to Ares (2012), then reverse transcribed using ProtoScript II with oligo-dT primers according to manufacturer’s instructions. cDNA product was treated with RNase H and RNase A, purified in a Zymo 5µg column, and amplified for 6 cycles with Q5 polymerase using primers RM512 and RM546 from Muller et al. (2020). This reaction was purified and amplified with distinct NEBNext dual index primer sets as above.

#### Computational analysis

RNA and DNA barcode reads were trimmed with cutadapt and quantified with custom scripts available in the Github repository. Barcodes with at least 24 reads in each DNA library were kept for analysis. Pre-vs post-induction RNA and DNA barcode abundance in two replicate turbidostat runs were analyzed with the ‘mrpa’ R package (Myint et al. 2019) to compute log fold changes of each barcode upon induction for the fast and slow reporter experiments.

### mRNA quantification by qRT-PCR

All qRT-PCR samples were prepared from frozen pellets from 1mL mid-log phase yeast culture. Total RNA was extracted using one-step hot formamide extraction, according to Shedlovskiy, et al. (2017). In brief, 25uL of a solution of 98% formamide and 10mM EDTA was used to resuspend 0.5 OD_600_ units of yeast and heated at 70°C for 10 min before being centrifuged at 21x *g* for 2 minutes. Supernatant containing the RNA was collected. RNA concentration was measured on a NanoDrop and 50µg RNA was purified using an RNA Clean & Concentrator-25 Kit (Zymo, R1017), with on-column DNaseI digestion, as per manufacturer’s instructions. RNA was eluted in 50µL of nuclease-free water and measured on a NanoDrop. 1µg of RNA was reverse transcribed with oligo(dT)_16_ primers and SuperScript IV (Invitrogen, 18090010) according to the manufacturer’s instructions. One-tenth of this reaction (2µL) was then subjected to qPCR with a DyNAmo HS SYBR Green qPCR Kit (Thermo Scientific, F410L) according to manufacturer instructions on a CFX96 Touch Real Time qPCR Detection System (Bio-Rad, 1845096 with 1851196). For each RT reaction, two sets of qPCR reactions were performed; one with primers specific to mCherry and one with primers specific to the citrine variant being probed, with each set consisting of three technical replicates. qPCR data were analyzed with the ΔΔCq method using custom R scripts (available on Github). The signal from each citrine variant was normalized to mCherry signal from the same biological sample.

### Polysome profiles

#### Yeast vacuum filtration and cryogrinding

Overnight cultures of the six citrine yeast strains were back-diluted so that each strain was at an OD of 0.05, for a combined OD of 0.1 in 125mL YEPD. Two mixed cultures were prepared, each consisting of different biological replicates of each citrine strain. Mixed cultures were shaken at 300 rpm at 30°C until they reached mid-log phase, at which point they were vacuum filtered using a sintered glass filter and 0.45 uM nitrocellulose membranes, then flash frozen in a falcon tube filled with LN2.

LN2 level was adjusted so that the yeast pellet was covered, then 2 mL of lysis buffer (10 mM Tris pH 7.0, 10 mM Tris pH 8.0, 150 mM NaCl, 5mM MgCl2, 1 mM DTT, 0.1mg/mL cycloheximide, 0.01% Triton-X, 0.0024 U/uL Turbo DNase) was added dropwise into sample. After washing, drying, and cooling the chamber and ball bearing of the Mixer Mill 400 (Retsch, 20.715.0001) in LN2, frozen yeast and lysis buffer was added to the chambers and shaken at 15 1/s frequency for 3 minute intervals 5 times. This cryoground lysate was then thawed, clarified via centrifugation, and its total RNA concentration was quantified using a Quant-It kit (ThermoFisher, Q33140).

#### Polysome fractionation and RNA extraction

To separate the polysomes, a standard polysome fractionation procedure was followed. Briefly, 10% sucrose (10% sucrose, 10 mM Tris pH 7.0, 10 mM Tris pH 8.0, 150 mM NaCl, 5mM MgCl2, 0.1mg/mL cycloheximide, 0.002 U/uL SuperAseIN) and 50% sucrose (50% sucrose, 10 mM Tris pH 7.0, 10 mM Tris pH 8.0, 150 mM NaCl, 5mM MgCl2, 0.1mg/mL cycloheximide, 0.002 U/uL SuperAseIN) solutions were prepared, then 8mL of 50% sucrose was layered under 6mL of 10% sucrose in Seton Open-Top Polyclear Centrifuge Tubes (Seton, 7030). Gradients were established using Biocomp Instruments’ Piston Gradient Fractionator (made up of part no.: 153, 105-914A-IR, and F-1-260). 200uL of clarified lysate was layered on top of these gradients and spun in a Beckman ultracentrifuge with a SW41 rotor at 36,000 rpm for 3 hours at 4°C. These gradients were then fractionated using the Piston Gradient Fractionator. Fractions were collected based on the polysome traces generated by the UV monitor, and subsequently flash frozen in LN2.

A constant amount of total RNA isolated from HEK293 (1uL of 500 ng/uL) was then spiked into each fraction and mixed. 500uL of each spiked-in fraction was aliquoted, ethanol precipitated overnight, and treated with DNaseI according to manufacturer’s instructions. Acid phenol chloroform extraction was then performed according to Ares (2012) to isolate the total RNA of each sample, and samples were ethanol precipitated once more and resuspended in water.

#### RNA-seq library preparation and analysis

These samples were sent to UC Berkeley’s Functional Genomics Laboratory for quantification and quality check via bioanalyzer, poly-A selection to enrich for mRNA, and Illumina library preparation. Samples were pooled in equimolar ratios, and Illumina sequencing was performed with the NovaSeq 6000 in an SP flow cell, with 100 single-read sequencing cycles.

To quantify citrine mRNAs in each fraction, RNA-seq reads were aligned with bowtie2 to a custom yeast transcriptome file containing all yeast genes plus the six citrine sequences. To calculate normalization from human mRNA spike-ins, reads were also aligned to the human transcriptome (allowing one alignment per read to avoid double-counting reads). The number of reads matching the fast and slow reporter in each polysome fraction was divided by the total number of human reads in that fraction. Analysis code is available on github.

### Translation simulations

We carried out stochastic simulations of translation that modeled the translation of an mRNA, followed by its eventual degradation, using a totally asymmetric simple exclusion process (TASEP) (MacDonald and Gibbs 1969; Zia et al. 2011). This simulation tracked the positions of ribosomes on a model transcript using the following elementary reactions:

- ***Ribosome loading:*** the recruitment of a small ribosomal subunit and scanning for the start codon. Loading is blocked when the mRNA is already occupied by another pre-initiation complex in the joining state (see below), or by another ribosome within 10 codons of the start codon.
- ***Subunit joining:*** the joining of a large and a small ribosomal subunit at the start codon. Joining was modeled with 3 independent sub-steps; upon completion of the final joining substep, the ribosome transitioned into an elongating state on the first codon.
- ***Elongation:*** translocation of a ribosome one codon along the mRNA. Elongation is blocked by another ribosome within 10 codons of the destination codon. Elongation on the stop codon represents translation termination, which immediately removes the ribosome and releases a completed protein.
- ***mRNA degradation:*** the transcript is instantaneously degraded and no protein is produced by ribosomes engaged in elongation.

We simulated initiation rates between 0.1 and 0.01 s^-1^, corresponding to an average interval of 10 to 100 seconds between initiation events. We chose 21 different values for the overall initiation rate, spaced logarithmically across this interval. The ribosome loading rate was set to twice the overall initiation rate, and the rate of each sub-step of the joining process was set to six times the overall initiation rate.

We determined per-position elongation rates by multiplying a “base” elongation rate of 6 s^-1^ by a per-position scaling factor determined from the iχnos model. The iχnos model predicts the time the ribosome spends on each position of a coding sequence as a function of the identity of the codon being decoded in the ribosomal A site as well as the adjacent codons in the P and E sites and neighboring sequence. The total elongation time modeled by iχnos for the slow reporter was 1.7 times longer than the total elongation time for the fast reporter.

We used an 0.0011 s^-1^ mRNA degradation rate for the fast reporter, corresponding to a 15 minute lifetime. This value is close to the half-lives measured for *PGK1* and *ADH1* transcripts, which were used to create the 5ʹ and 3ʹ UTRs of the reporters. We accelerated mRNA degradation of the slow reporter 1.54-fold to match the measured steady-state abundance ratio between the slow and fast reporters, under the assumption that their transcription rates were identical.

We carried out 16,384 replicate simulations for each parameter set using the Gillespie algorithm (Gillespie 1976) and recorded mRNA lifespan, protein production, and the frequency of “failed” reactions that were blocked by conflicts with other ribosomes on the same mRNA.

## Supplemental Figures

**Figure S1 (corresponds to Fig. 2 and 4):**
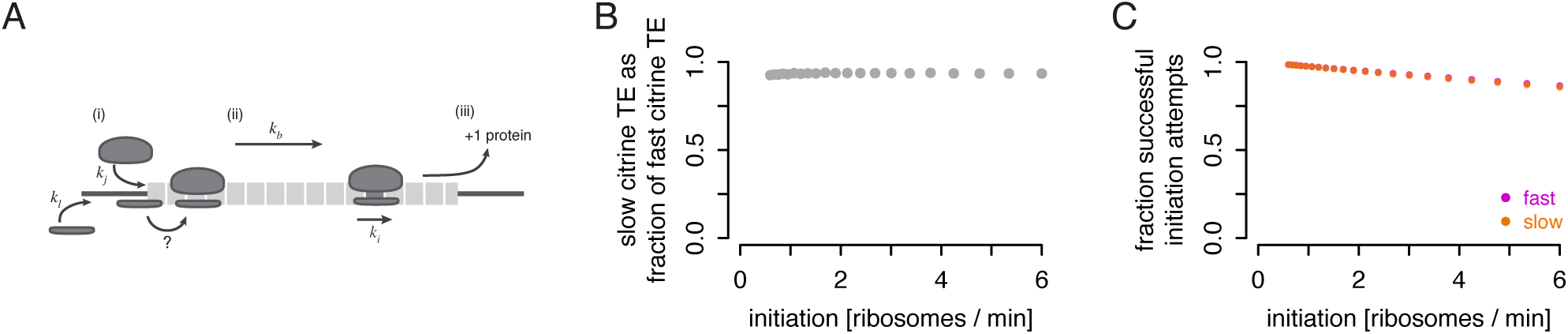
Schematic of the basic Totally Asymmetric Simple Exclusion Process (TASEP) model simulating translation of our citrine sequences. There are four steps: (i) pre-initiation complexes will load on to the 5′ end of the mRNA and scan for the start codon with loading rate k_l_, as long as they are not blocked by another ribosome overlapping the start codon. (ii) Ribosomal subunits will then join to form an elongation complex with three independent steps, each at rate k_j_. (iii) As long as there is no ribosome blocking the one upstream, ribosomes will move to the next codon with rate k_i_, which is equal to the base elongation rate, k_b_ multiplied with the elongation rate factor at every position i. This factor i is relative, and is calculated as (1/occupancy rates) from the iXnos algorithm. (iv) Ribosomes terminate when they reach the end of the coding sequence, and one protein is counted. b) The difference in the ratio of slow to fast simulated translation efficiency (protein/mRNA*time) from the TASEP model is very small across a range of realistic initiation rates. c) Percentage of initiation events that succeed without interference over a range of initiation rates in TASEP simulations. The impact of interference does not differ substantially between fast and slow reporters.

**Figure S2 (corresponds to Fig. 5):**
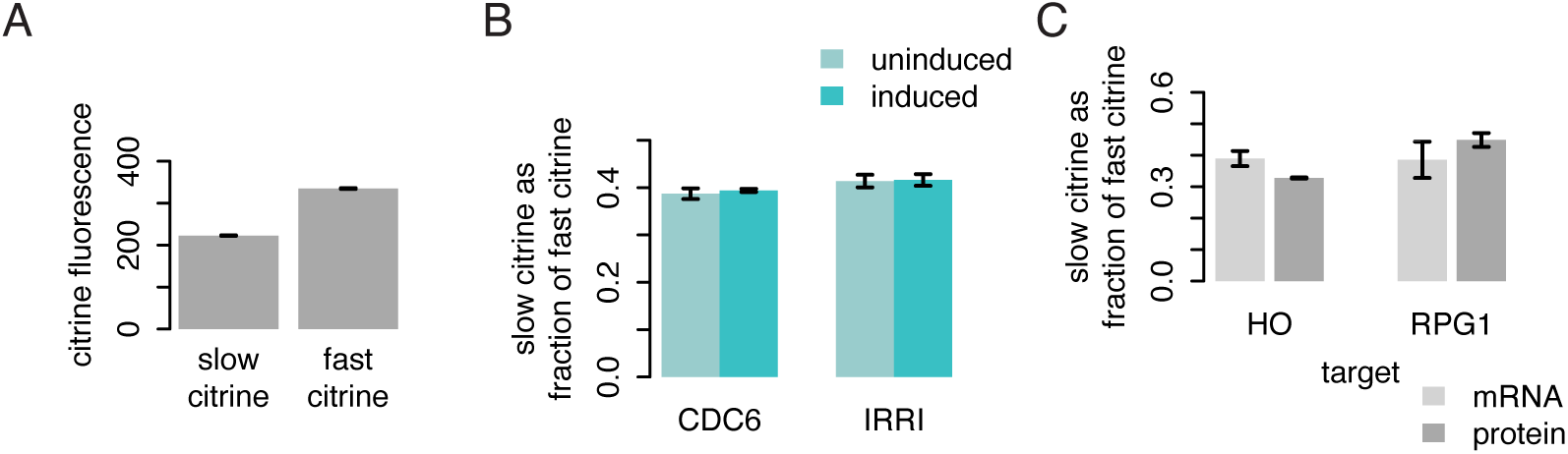
a) Fluorescence of the ZEM TF–citrine fusions used in the CiBER-seq screen was measured with flow cytometry. The medians of ∼15,000 events are plotted after background correction, showing a difference in fluorescence of slow vs fast fusions. b) Ratio of normalized fluorescence of the slow citrine reporter to normalized fluorescence of the fast citrine reporter, as in figure 5C, in strains with CRISPRi guides against genes that are not involved in translation but whose knockdown causes large growth defects (McGlincy et al., 2021). The strains exhibit no differential effect on citrine fluorescence. c) Relative abundance of mRNA from slow vs fast citrine reporters was similar between CRISPRi knockdown of RPG1 and the HO locus control. mRNA abundance (light gray) was determined by RT-qPCR from three isolates, normalized to mCherry mRNA. Data showing a change in fluorescence ratios (dark gray) are repeated from figure 5C for comparison.

## Notes

### Summary of Updates

New experimental data and analyses; substantial revisions throughout the manuscript

